# Estimating conditional cell population associations across tumor density trajectory in three dimensional tissue images

**DOI:** 10.64898/2026.04.16.719017

**Authors:** Sagnik Bhadury, Arvind Rao

## Abstract

Registered serial-section imaging preserves a depth coordinate that a single planar image cannot represent. We introduce ISPat-3D (Informed Spatially Aware Patterns in Three Dimensions), a framework that combines anisotropic Gaussian-process adjustment with multi-study factor analysis to separate smooth spatial variation from covariance shared across tumor-density zones and covariance specific to individual zones. Precision matrices summarize the adjusted structure as signed partial correlations, interpreted as conditional co-location or exclusion rather than physical contact, signaling, or causation. We analyze conditional associations among cell-type neighborhood-density variables across five relative tumor-burden strata in one colorectal-cancer CyCIF specimen and one HER2-positive breast-carcinoma IMC specimen. The workflow adjusts modeled smooth spatial variation with an anisotropic three-coordinate Gaussian process, fits shared and zone-specific factor covariance to the residuals, and converts each full zone covariance to signed partial correlations. To make the reported cell budgets feasible, the analysis uses a 15-neighbor Vecchia GP likelihood on spatially balanced anchors and a Gaussian-likelihood fit of the same factor covariance form from full residual covariance summaries. Controlled simulations, a matched section-wise planar comparison, and direct graphical-lasso sensitivity check the estimator. In CRC, positive T-cell and macrophage–stroma density associations show relatively high agreement across three sampling budgets. Breast pair signs are much less stable across budgets and remain exploratory, even though graphical lasso agrees on the same GP-adjusted residuals. A GP separates patterns by assumed spatial scale; a smooth field may contain meaningful biology and residual associations may contain noise or unmeasured effects. The two single-specimen analyses identify biologically plausible, zone-resolved conditional associations for replication, without establishing cellular contact, mechanism, or cohort-level reproducibility. Code release: https://github.com/sagnikbhadury/ISPAT-3D.

## Introduction

The tumor microenvironment (TME) is a spatially organized mixture of malignant, immune, vascular, and stromal cells whose local arrangement can influence immune access, tumor progression, and treatment response. Their distributions vary with tumor density, distance from an invasive margin, and local immune infiltration^1,2^.

Multiplexed imaging studies have consequently shifted attention from whole-tissue cell abundance toward cellular neighborhoods and tissue compartments; in colorectal cancer, for example, coordinated neighborhoods at the invasive front distinguish immune-active from immune-suppressive organization^3^. A whole-tissue average can conceal such localized associations, whereas a single pairwise proximity statistic can confound a direct relationship with the shared tendency of several cell types to occupy the same compartment.

Multiplexed platforms such as cyclic immunofluorescence (CyCIF), imaging mass cytometry (IMC), multiplexed ion-beam imaging, and CODEX quantify many proteins while retaining cell locations^4–7^. Most analyses nevertheless operate on one tissue section at a time. A thin section can split a connected structure, miss a rare focal niche, or make two populations appear adjacent even when their three-dimensional trajectories diverge. Serial-section imaging supplies a physical depth coordinate, allowing vessels, tumor fronts, immune aggregates, and stromal boundaries to be followed through a volume^8–10^. Recent reconstructions show that three-dimensional context can resolve connected tumor subclones and immune neighborhoods across sections^11^; quantitative 3D histology further demonstrates that immune-hot and immune-cold regions may alternate over tens of micrometres around the same precursor lesion^12^. These findings motivate statistical summaries that use the registered volume rather than treating serial sections as exchangeable replicates.

The biological question is how the network of local cell-type densities changes across heterogeneous tumor regions. We first turn the locations of each annotated cell type into a smooth local-density surface by kernel density estimation (KDE), so that every sampled cell location has a vector of neighborhood densities. We then order locations by a tumor-burden measure and divide them into five relative zones, from Very Low to Very High. In CRC, that measure is the tumor-cell KDE estimated within each section; in breast carcinoma, it is the measured pan-cytokeratin (panCK) intensity at each cell, used as an epithelial-burden proxy. The same cell-type KDE variables are analyzed within each zone. Comparing their conditional-association networks across these ordered zones addresses whether relationships among cell populations differ between tumor-sparse and tumor-rich parts of the specimen.

The analytical challenge is not simply to calculate distances in three dimensions. Broad gradients in tumor burden, stromal content, staining, or tissue geometry can raise several cell-type density surfaces together and thereby create marginal co-occurrence without a corresponding conditional association. Conversely, fitting each small region independently discards information that could stabilize estimation when cell types are sparse. A useful model for registered tumor volumes should therefore distinguish smooth spatial background from cross-variable dependence, condition each association on the remaining measured cell types, and borrow information across biologically ordered regions while allowing genuine regional departures.

Existing methods address important parts of this problem but target different estimands. Nearest-neighbor summaries, neighborhood clustering, inhomogeneous point-process methods, SpaceANOVA, and Ripley’s *K* characterize proximity or pairwise co-occurrence; these constructions are not intrinsically two-dimensional, and an explicit three-dimensional *K* estimator is available in the spatstat software^13–15^. Spatial cellular graphs and graph-learning models capture disease-relevant local motifs^16^, while SpatialDE and related Gaussian-process models test or decompose spatial variation in molecular measurements^17–19^. CellPhoneDB and COMMOT instead use expression and ligand–receptor information to infer candidate communication pathways^20,21^. More recent spatially varying graphical models estimate continuously varying conditional edges with posterior uncertainty, but address two-dimensional tissue images rather than registered serial-section volumes partitioned along an ordered tumor-density axis^22^. Thus proximity, communication, spatial-variance, and conditional-network methods are complementary rather than interchangeable.

The remaining motivation is to represent both recurrence and heterogeneity. Tumor-density zones are related microenvironments, not unrelated datasets: some covariance may recur throughout the specimen, whereas other structure may emerge only toward the tumor-dense core or tumor-sparse periphery. Estimating every zone separately can be unstable and obscures what is shared; pooling all zones erases the very heterogeneity of interest. Multi-study factor analysis provides a parsimonious compromise by decomposing covariance into shared and study-specific low-rank components^23,24^, with variational algorithms making the decomposition practical for repeated fits^25^. In this manuscript, the “studies” are ordered tumor-density zones, and the resulting factor components are converted to partial-correlation graphs for descriptive conditional-association analysis.

ISPat-3D builds on our earlier two-dimensional ISPAT framework^26^. Its contribution is the integration of registered (*x, y, z*) coordinates, an anisotropic GP with a distinct axial length scale, ordered tumor-density strata propagated across sections, and an MSFA decomposition into shared and zone-varying covariance. We borrow GP and MSFA methodology rather than propose a new GP or factor model. A GP is a distribution over smooth spatial functions; here it estimates broad variation that could otherwise induce apparent co-occurrence. A loading measures how strongly a cell-type density follows a latent factor, with one loading matrix shared across zones and another specific to each zone. After covariance estimation, a network visualizes partial correlations: an edge means residual conditional association after accounting for the other measured cell types, not physical contact, ligand–receptor signaling, or causation.

The method has five conceptual steps (Figure 1): start with a registered 3D cell map; divide the volume into ordered, non-overlapping tumor-density zones; attenuate modeled smooth spatial structure with an anisotropic GP; use MSFA to separate shared from zone-varying covariance; and convert each full fitted zone covariance to a partial-correlation graph. This ordering separates the biological question from the statistical machinery. Figure 2 shows the data-level CRC pipeline, from the registered volume and tumor-density zones to illustrative shared and zone-specific network outputs.

**Fig 1.**
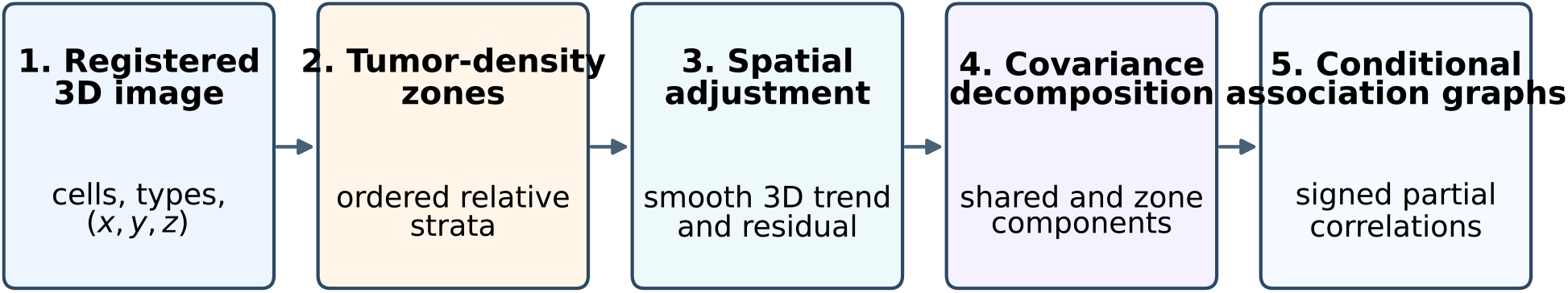
Conceptual ISPAT3D study workflow connecting the registered image to zone-specific conditional-association graphs in five stages. Edges represent fitted conditional density associations rather than physical contact or avoidance.

**Fig 2.**
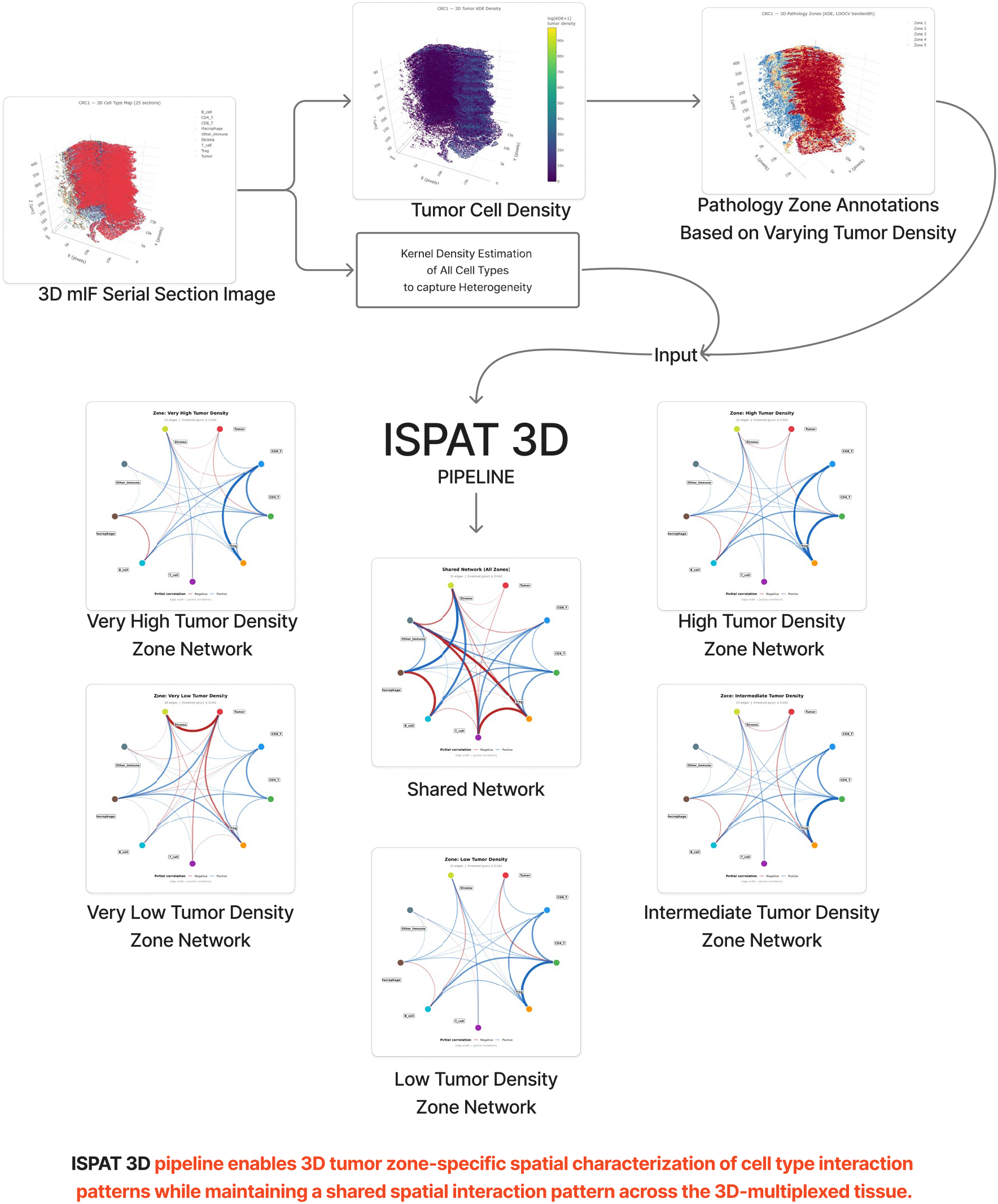
Data-level ISPAT3D pipeline. The registered CRC volume yields a tumor-cell density surface, relative tumor-density zones, and cell-type-specific neighborhood-density inputs. The shared and zone-specific network drawings illustrate the output structure; Figures 5 and 6 present the fitted CRC associations. An edge denotes a statistical conditional density association, not direct cell contact.

The main interpretive assumption is that sufficiently smooth variation is a background field to be adjusted before analyzing local residual association. A smooth field may also contain important biology, including immune exclusion, stromal boundaries, and vascular organization; residuals may contain noise, annotation errors, and unmeasured spatial variation. We therefore compare spatial scales in a controlled simulation, display effect sizes and sign stability for every pair, fit a matched section-wise 2D comparator, and perform a direct graphical-lasso sensitivity analysis. These checks delimit the interpretation of edges; they cannot settle the compartment-versus-interaction question in a single image^27^.

The rest of the manuscript is organized as follows. Section 2 presents simulations of covariance and magnitude-threshold recovery, computation time, and GP behavior under shared spatial confounding, followed by colorectal and breast-carcinoma analyses, matched section-wise ISPAT-2D comparisons, and direct residual-precision sensitivity. Section 3 discusses biological interpretation, methodological scope, reproducibility, limitations, and future work. Section 4 details the simulation designs and ISPat-3D formulation, including tumor-burden zoning, cell-type KDE inputs, anisotropic GP adjustment, shared and zone-specific covariance estimation, partial-correlation networks, stability checks, the planar comparator, and optional reference-informed regularization. Appendix A expands the zone-specific network and stability interpretation; Appendix B documents graphical-lasso sensitivity; and Appendices C and D describe specimen processing and neighborhood-density construction.

## Results

The five labels Very Low through Very High are specimen-specific, relative tumor-burden strata: CRC uses tumor-KDE quintiles and breast uses panCK-intensity quintiles. Tied breast scores make the observed zone counts unequal. The labels are not discrete biological states. An edge is a signed partial correlation between two adjusted cell-type density variables, conditional on the other measured variables. Its magnitude, agreement across sampling budgets, and agreement in section-block resamples are descriptive checks; none is a statistical-significance test.

### Simulation study

We analyzed the generated samples with GP and covariance-likelihood estimators. The design included three zones, *G* ∈ {15, 20, 25} variables, six balanced or moderately imbalanced zone-size configurations of 250–300 observations per zone, Matérn-3/2 and RBF generating kernels, and five replications per setting (180 data sets). Shared and zone-specific factor covariance plus diagonal uniqueness generated the nonspatial signal; independent anisotropic fields generated a spatial nuisance. The RV coefficient^28^ measures similarity between positive-semidefinite matrices. Truth and fitted zone matrices were compared by the RV coefficient, and entries of absolute covariance exceeding 0.1 were compared as a descriptive magnitude-recovery measure. Because the factor-generated truth is dense, the latter is not an inferential edge-selection or false-discovery-rate guarantee.

Across the 180 fits, the mean RV was 0.887 for the shared factor covariance and 0.966, 0.963, and 0.964 for the three full zone covariances (Figure 3). The mean sensitivity under the 0.1 magnitude rule was 0.866 for the shared component and 0.934–0.938 for zone matrices; the corresponding descriptive false-discovery proportions were 0.033 and 0.023–0.027 (Figure 3). These summaries describe the specified estimator.

**Fig 3.**
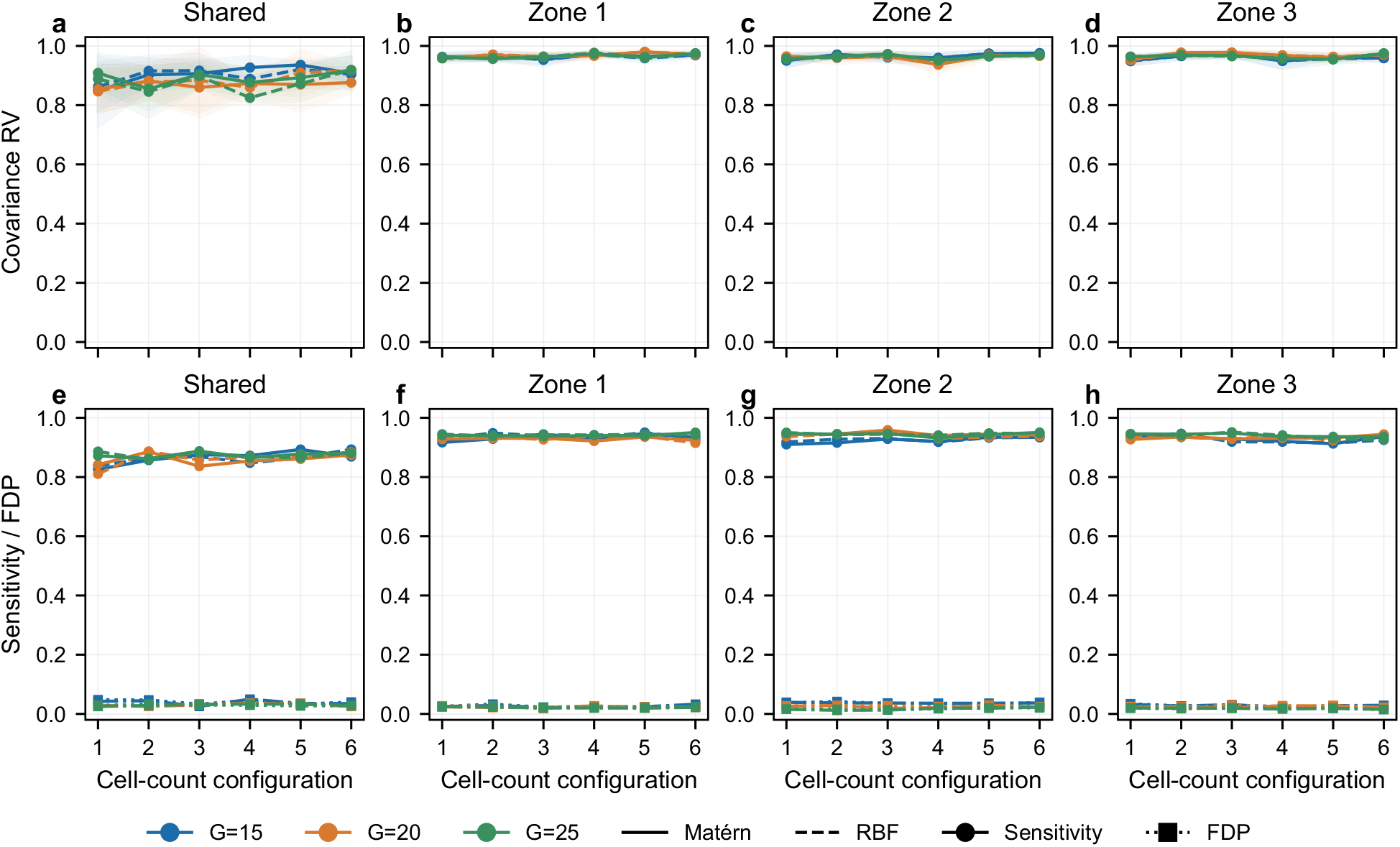
Simulation recovery across all *G* = 15, 20, and 25 settings in one faceted figure. Panels (**a**–**d**) show covariance RV recovery for the shared and three full zone covariances. Panels (**e**–**h**) show sensitivity and descriptive false-discovery proportion under the 0.1 magnitude rule. Shaded RV bands show replicate standard deviations; points summarize five replications per setting. These magnitude summaries are not inferential edge-wise error control.

Observed GP-stage wall time averaged 9.34 seconds (range 6.06–12.94) for these small simulation fits, and a separately timed MSFA covariance fit averaged 0.58 seconds (Figure 4). GP runs shared the machine with other concurrent jobs; these timings describe the observed environment and should not be extrapolated to whole-slide workloads.

**Fig 4.**
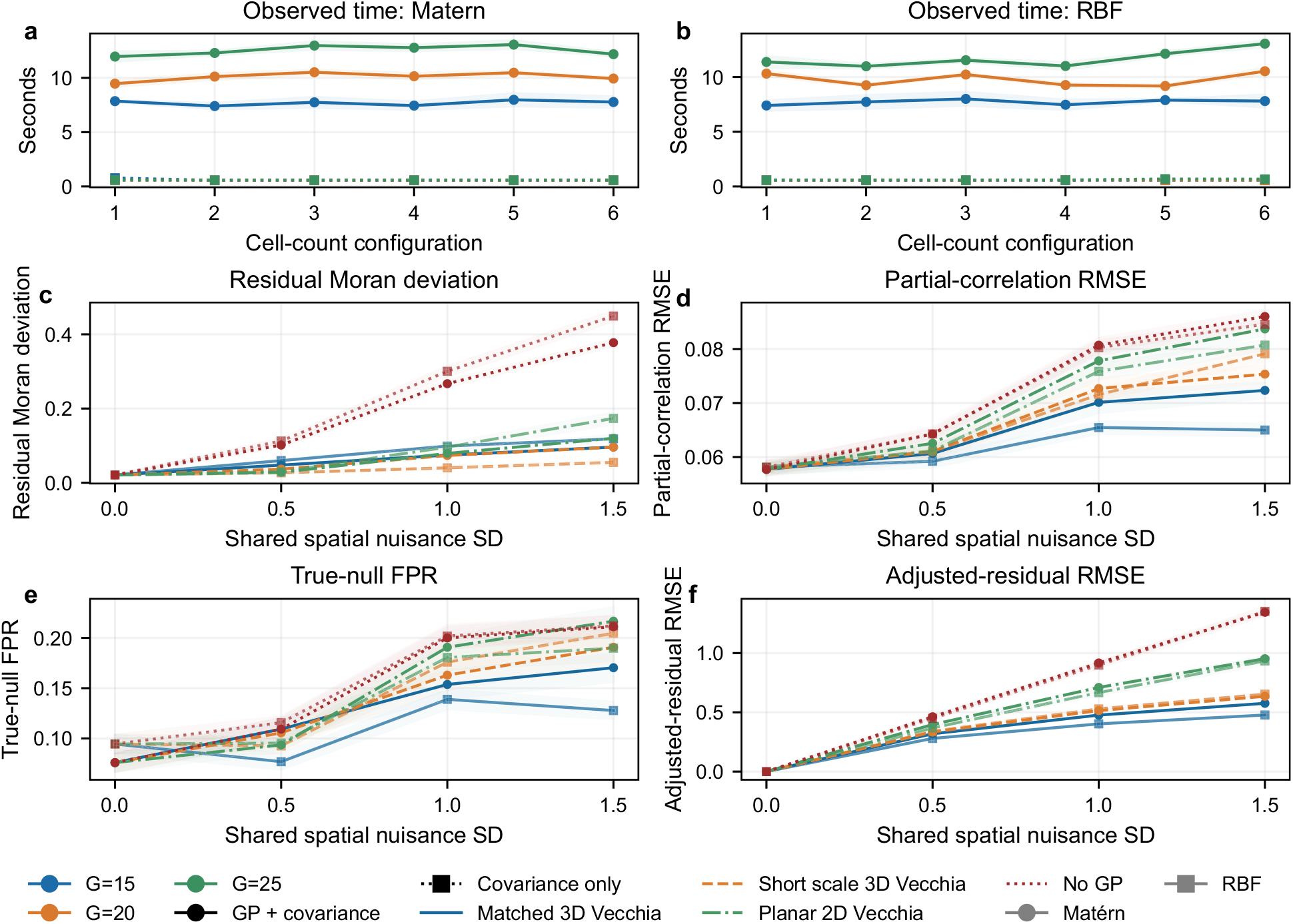
Computation and focused Vecchia GP validation. Panels (**a**,**b**) show observed timing for Mat’ern and RBF simulations across all *G* values. Panels (**c**–**f**) compare matched 3D, short-scale 3D, matched planar, and no-GP adjustment for residual spatial autocorrelation, partial-correlation RMSE, true-null false-positive rate, and adjusted-residual RMSE. Shaded bands summarize timing replicate standard deviations or validation standard errors, respectively. All validation arms use identical generated data within each replicate.

A focused component experiment used 12 variables and 300 three-dimensional locations with a known sparse ring precision matrix. Two shared smooth fields were added at four nuisance strengths *σ*_*B*_ ∈ {0, 0.5, 1.0, 1.5} under two kernels; each setting had 20 replicates. The same simulated realization was processed by a GP matched to the generating 3D spatial scale, a GP with half that scale, a matched planar (*x, y*) GP, and mean-centering without a GP. Known covariance parameters isolated the residualization step; the Vecchia approximation was used for the GP arms. At zero nuisance all four arms reduced to mean-centering. At *σ*_*B*_ = 1.5 with Matérn fields, the matched 3D arm had mean absolute residual Moran deviation 0.096 and partial-correlation RMSE 0.072, versus 0.120 and 0.084 for the planar GP and 0.378 and 0.086 without GP. Under RBF fields these values were 0.118 and 0.065 for matched 3D, 0.173 and 0.081 for planar, and 0.449 and 0.085 without GP. The short-scale 3D arm sometimes removed more residual spatial autocorrelation while worsening graph recovery, showing sensitivity to the assumed spatial scale. Figure 4 also gives true-null edge false-positive rates under the descriptive |*ρ*| > 0.1 rule. This controlled experiment supports using depth when the nuisance actually varies over registered *z*; it cannot prove that smooth variation in a real tumor is biologically irrelevant.

### Colorectal cancer application

We applied ISPat-3D to the colorectal cancer (CRC) cyclic immunofluorescence (CyCIF) dataset released by^9^. The dataset comprises a single specimen subjected to full 3D serial section imaging (CRC1). Since ISPat-3D requires a true 3D spatial structure – namely, consistent global XY coordinates across registered serial sections and a physical Z axis – CRC1 satisfies the structural requirements of the method.

CRC1 is a poorly differentiated stage IIIB BRAF^V600E^ colorectal adenocarcinoma with high microsatellite instability and complex histomorphology. The specimen was serially sectioned into 106 FFPE slices of 4 *µ*m thickness, from which 25 CyCIF sections were selected at non-consecutive physical indices to maximize the total reconstructed depth along the Z-axis while skipping interleaved H&E sections. Each section was assigned a physical depth coordinate *Z*_*µ*m_ = *s* × 4, where *s* is the section index, yielding a total reconstructed depth spanning 8 to 424 *µ* m. Whole-slide CyCIF imaging was performed using 102 antibodies covering epithelial, immune, and stromal lineage markers as well as markers of cell cycle state, signaling activity, and immune checkpoint expression^9^. CyCIF image tiles and imaging cycles were stitched and registered, and cells were segmented, using MCMI-CRO^29^. The CRC atlas provided tissue-level XY coordinates (*X*_*s*_, *Y*_*s*_) registered across all 25 physical sections^9^.

The CRC1 CyCIF volume contains 9,431,098 processed KDE rows across 25 registered sections. Nine celltype density variables were analyzed. Spatially balanced sampling provided 50,000, 100,000, and 150,000 cells *per zone* in the five relative tumor-density quintiles; the largest fit therefore used 750,000 selected cells in total. The selected source row identifiers, GP residuals, fit metadata, and factor covariance outputs are saved for every tier. Each cell-type–zone GP was fit with a Matérn anisotropic three-coordinate Vecchia likelihood on spatially balanced anchors and used to predict the spatial field at all selected cells. The full residual covariance summaries and actual row counts then informed the shared-plus-zone factor covariance fit.

Figure 5 shows all five circular zone-network views in one faceted figure, using the circular network layout and the fitted partial correlations. Figure 6 complements the thresholded topology view by displaying all 36 cell-type pairs across five zones, with signed partial-correlation magnitudes, agreement of the edge sign across the three budgets, and the proportion of 100 serial-section block resamples retaining that sign. In its upper panels, “Effect” is the signed partial correlation; “Budget sign” and “Block sign” are proportions of fits retaining the aggregate sign across the three budgets and 100 fixed-residual section-block resamples, respectively. Overall, 169 of 180 pair–zone entries (93.9%) kept the same sign across all three budgets; 87.2% reached at least 90% sign agreement in the fixed-residual block resamples. Positive CD4^+^ T–Treg partial correlations were 0.213, 0.273, 0.368, 0.357, and 0.296 from Very Low to Very High; CD8^+^ T–CD4^+^ T values were 0.348, 0.334, 0.247, 0.206, and 0.163. Macrophage–Stroma values were 0.221, 0.188, 0.157, 0.153, and 0.148. These are conditional density associations in this specimen and do not establish contact, immune suppression, or a causal relation.

**Fig 5.**
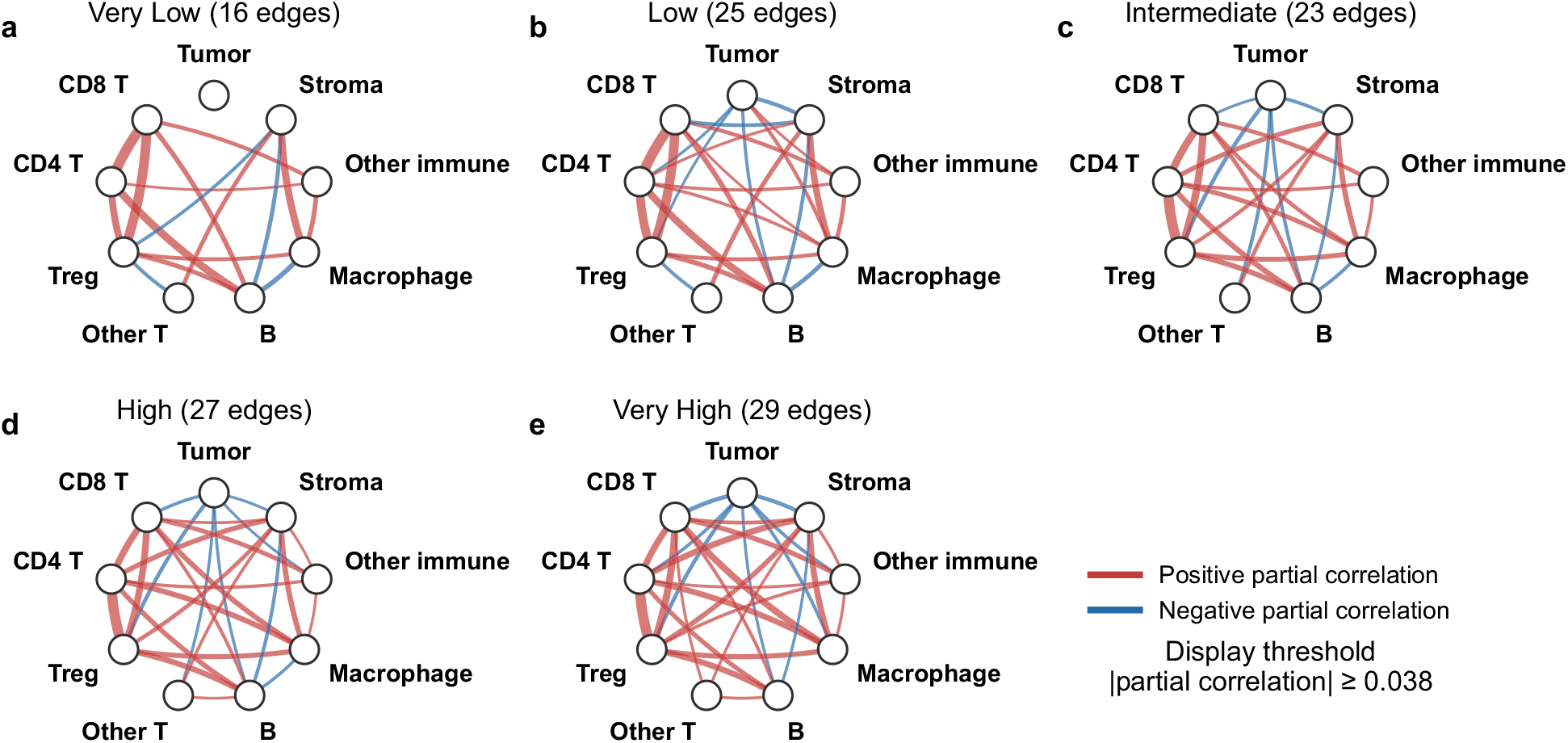
CRC zone-specific conditional-association networks in a circular layout. Panels (**a**–**e**) show the five relative tumor-density strata, derived from the fitted partial correlations. Red and blue edges denote positive and negative partial correlations; width scales with magnitude. The display threshold |*ρ*| ≥ 0.038 reduces crossings.

**Fig 6.**
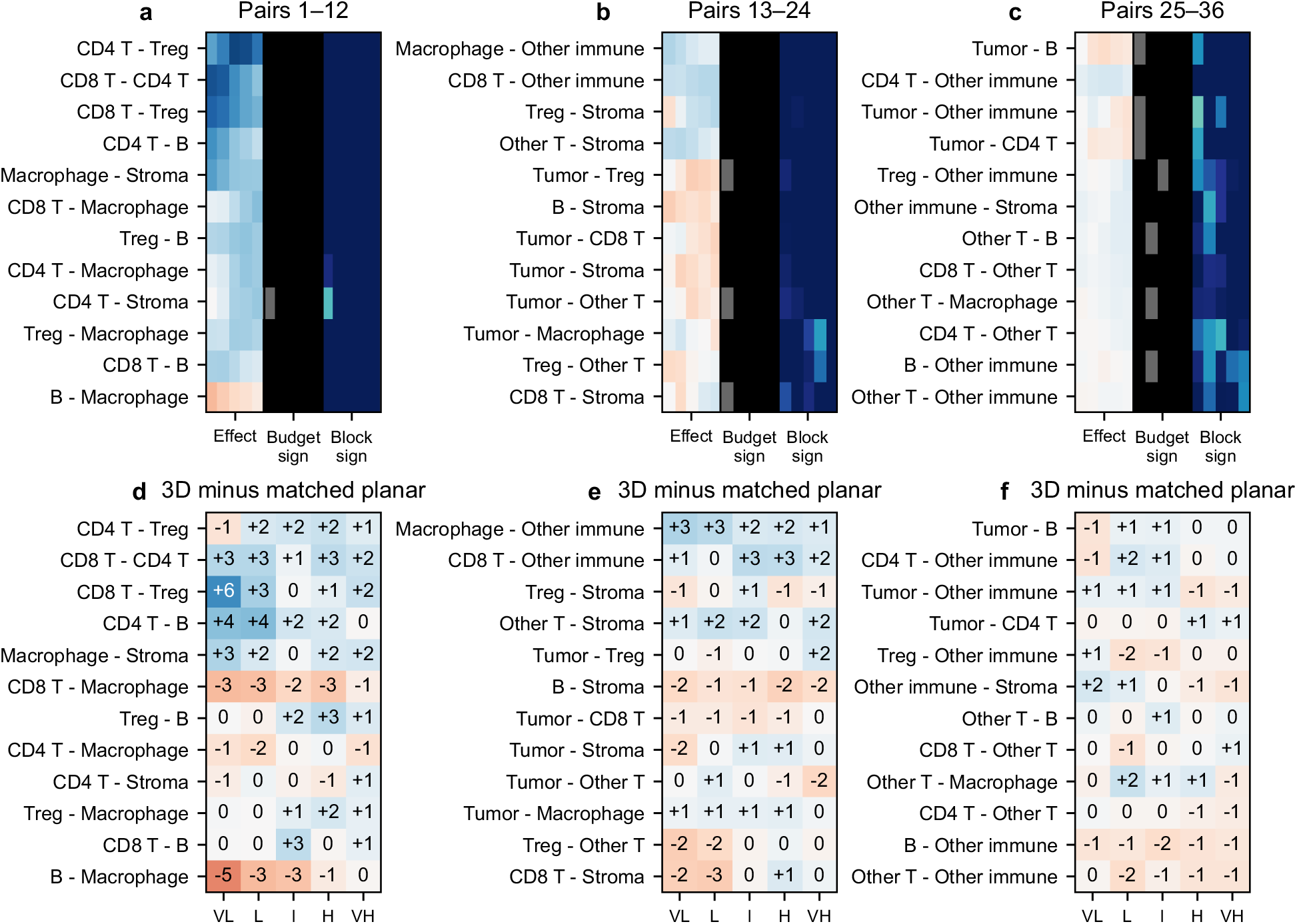
Complete CRC quantitative comparison. Panels (**a**–**c**) partition all 36 cell-type pairs across five zones. “Effect” is the signed fitted partial correlation for a pair and zone. “Budget sign” is the fraction of the three actual cell-sampling budgets whose edge sign matches the aggregate sign (1/3, 2/3, or 3/3). “Block sign” is the fraction of 100 serial-section block resamples retaining that sign with GP residuals held fixed. Panels (**d**–**f**) show corresponding 3D-minus-matched-section-wise-2D partial correlations; cell numbers are differences in hundredths (e.g., +6 denotes approximately +0.06). Signed effect colors span −0.4 to +0.4; darker budget and block-sign cells represent stronger agreement. Stability measures are descriptive, and the lower panels are pipeline contrasts rather than coordinate-specific effects.

*Zone-specific interaction patterns*. In the Very Low CRC zone, the leading positive fitted associations are CD8^+^ T–CD4^+^ T (0.348), CD8^+^ T–Treg (0.324), CD4^+^ T–B cell (0.243), and Macrophage–Stroma (0.221). The same T-cell module remains positive in Low, where CD8^+^ T–CD4^+^ T and CD8^+^ T–Treg are 0.334 and 0.301. In Intermediate and High, CD4^+^ T–Treg becomes the largest positive association (0.368 and 0.357); the corresponding Very High value is 0.296. This is a shift in relative fitted magnitude across density strata, not evidence that Treg function changes with tumor burden. CD8^+^ T–CD4^+^ T decreases from 0.348 in Very Low to 0.163 in Very High, while CD4^+^ T–B cell decreases from 0.243 to 0.098. All three sampling budgets preserve the sign of these selected pairs in every zone. The coordinated T-cell patterns are compatible with reported immune neighborhoods in CRC^3^ and local Treg–CD8^+^ T proximity^30^, but the present density associations do not resolve cellular contact or a mechanism.

Macrophage–Stroma is positive in all five CRC zones, declining from 0.221 in Very Low to 0.148 in Very High. This persistence is compatible with the stromal–myeloid organization described in the source CRC volume and independent spatial analyses of FAP^+^ fibroblasts and SPP1^+^ macrophages^9,31^. CD4^+^ T–Macrophage rises from 0.034 in Very Low to 0.155 in High and 0.141 in Very High. Tumor–Stroma remains weakly negative (−0.015 to −0.090), illustrating that the network contains both positive and negative conditional density associations. These are specimen-level patterns; the numerical changes do not by themselves identify a biological transition.

The matched planar comparison used exactly the central 100,000-per-zone CRC cell roster. Planar Matérn Vecchia GPs were fit within each section and zone using (*x, y*) coordinates, after which the same factor covariance and partial-correlation steps were applied. Figure 6 shows all signed 3D-minus-2D differences; the median absolute difference was 0.011 and the largest was 0.063 (CD8^+^ T–Treg in Very Low). These are differences between pooled volumetric GP adjustment and independent section-wise planar GP adjustment, followed by matched downstream fitting. They do not isolate the *z* coordinate alone as the cause of an application edge.

### Breast carcinoma application

We also applied ISPat-3D to the three-dimensional imaging mass cytometry (IMC) dataset released by^8^. The dataset comprises a single HER2-positive ductal breast carcinoma (BC) specimen subjected to full volumetric 3D IMC acquisition. Since ISPat-3D requires a true 3D spatial structure with consistent global XY coordinates across registered serial sections and a physical Z axis, this specimen satisfies the structural requirements of the method.

The tissue block was sectioned into 152 consecutive 2 *µ*m FFPE slices using an ultramicrotome with a diamond knife, following the protocol described in^8^. Each section was assigned a physical depth coordinate *Z*_*µ*m_ = *s s*, where *s* is the section index, yielding a total reconstructed depth of 304 *µ*m. IMC acquisition was performed using a panel of 26 metal-conjugated antibodies covering epithelial lineage markers (pan-cytokeratin, CK5, CK7, CK8/18, CK14, CK19, E/P-cadherin, HER2), immune cell markers (CD3, CD8a, CD20, CD45, CD68, CD138), stromal markers (SMA, Vimentin, Collagen I, vWF/CD31), proliferation and apoptosis markers (Ki-67, phospho-H3, pS6, cPARP/cCasp3), and structural markers (Histone H3, Ir193). Images from the overlapping area across all 152 sections were aligned, and single cells were segmented using a 3D watershed algorithm applied to the Ir193 nuclear channel, yielding globally consistent XY coordinates for each segmented cell that are directly comparable across all sections. Mean marker intensities per cell were computed over each single-cell 3D mask and exported as a cell data catalog. Metal isotope spillover between channels was corrected using the compensation matrix described in^8^.

The processed HER2-positive IMC breast specimen has 73,936 cells in total across 152 sections. The 25,000, 45,000, and 65,000 budgets are *total cells across the five zones*, allocated in proportion to their available cell counts. Twelve KDE variables were retained for reporting, but the luminal tumor variable was structurally zero and excluded from active factor fitting before being re-embedded as a zero-valued row and column. In the central 45,000-cell planar comparison, 30 cells in very small section–zone groups were mean-centered because fewer than ten cells were available for a stable within-group GP.

Figure 7 shows all five circular zone-network views in one faceted figure, using the circular network layout and the fitted partial correlations. Figure 8 provides the complete effect, stability, and matched-planar comparison for every pair among the 11 active variables across five zones (275 pair–zone entries), with magnitude and budget and section-block sign stability. Excluding the structurally zero luminal variable, 187 of 275 entries (68.0%) kept the same sign across all three budgets, and 171 of 275 (62.2%) reached 90% sign agreement under fixed-residual section-block resampling. HER2-positive tumor–basal tumor was negative in the Intermediate, High, and Very High zones (−0.290, −0.366, and −0.374), but was 0.042 in Low and showed weaker budget agreement there. CAF– Myoepithelial partial correlations were modest (0.032–0.098 across zones), across zones. Endothelial–CAF values were 0.039–0.093. We treat all these breast patterns as candidates for replication, particularly because annotation composition, section heterogeneity, and spatial-scale choice can affect them.

**Fig 7.**
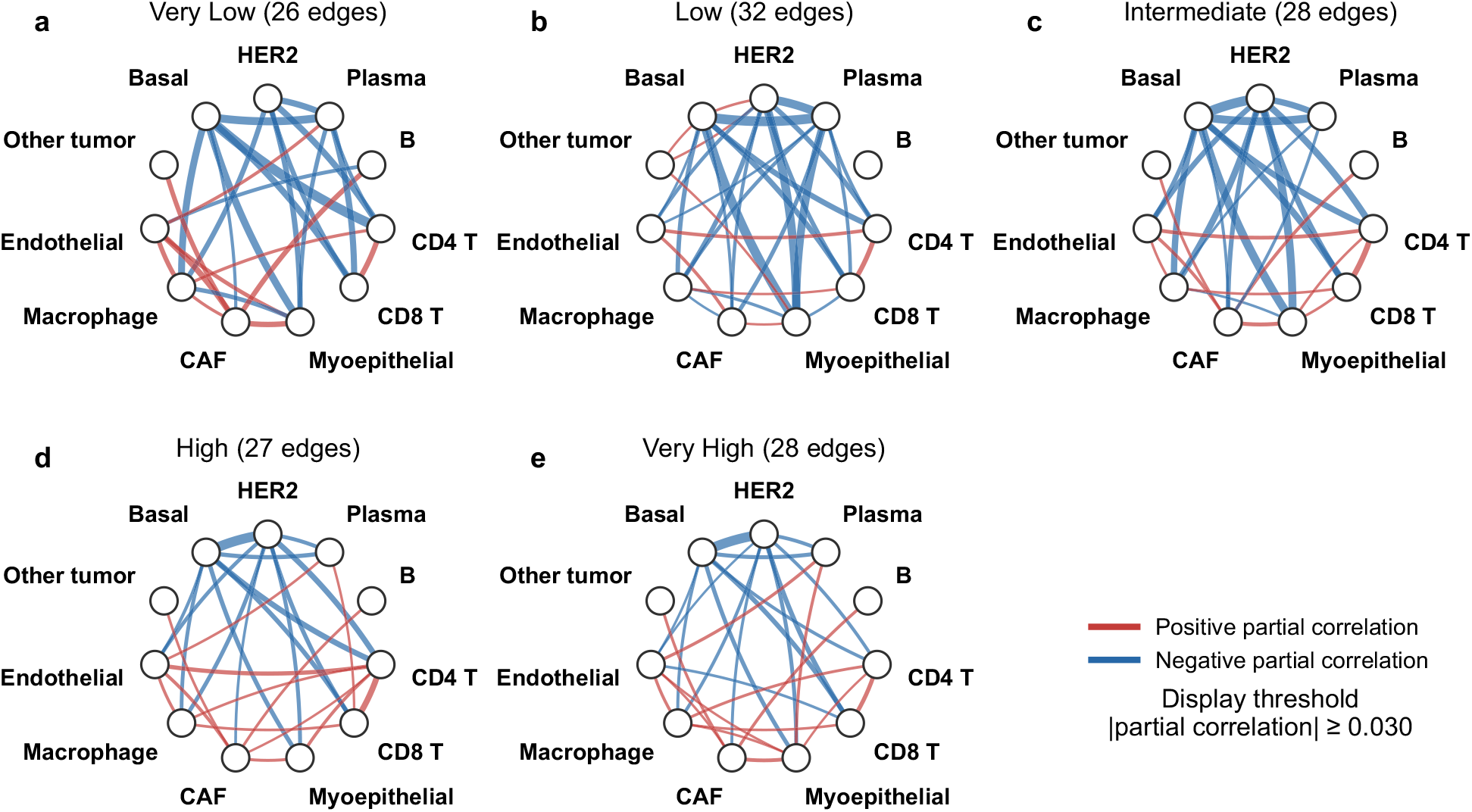
Breast zone-specific conditional-association networks in a circular layout. Panels (**a**–**e**) show the five relative panCK-burden strata for the 11 active variables, derived from the fitted partial correlations. Red and blue edges denote positive and negative partial correlations; width scales with magnitude. The display threshold |*ρ*| ≥ 0.030 reduces crossings.

**Fig 8.**
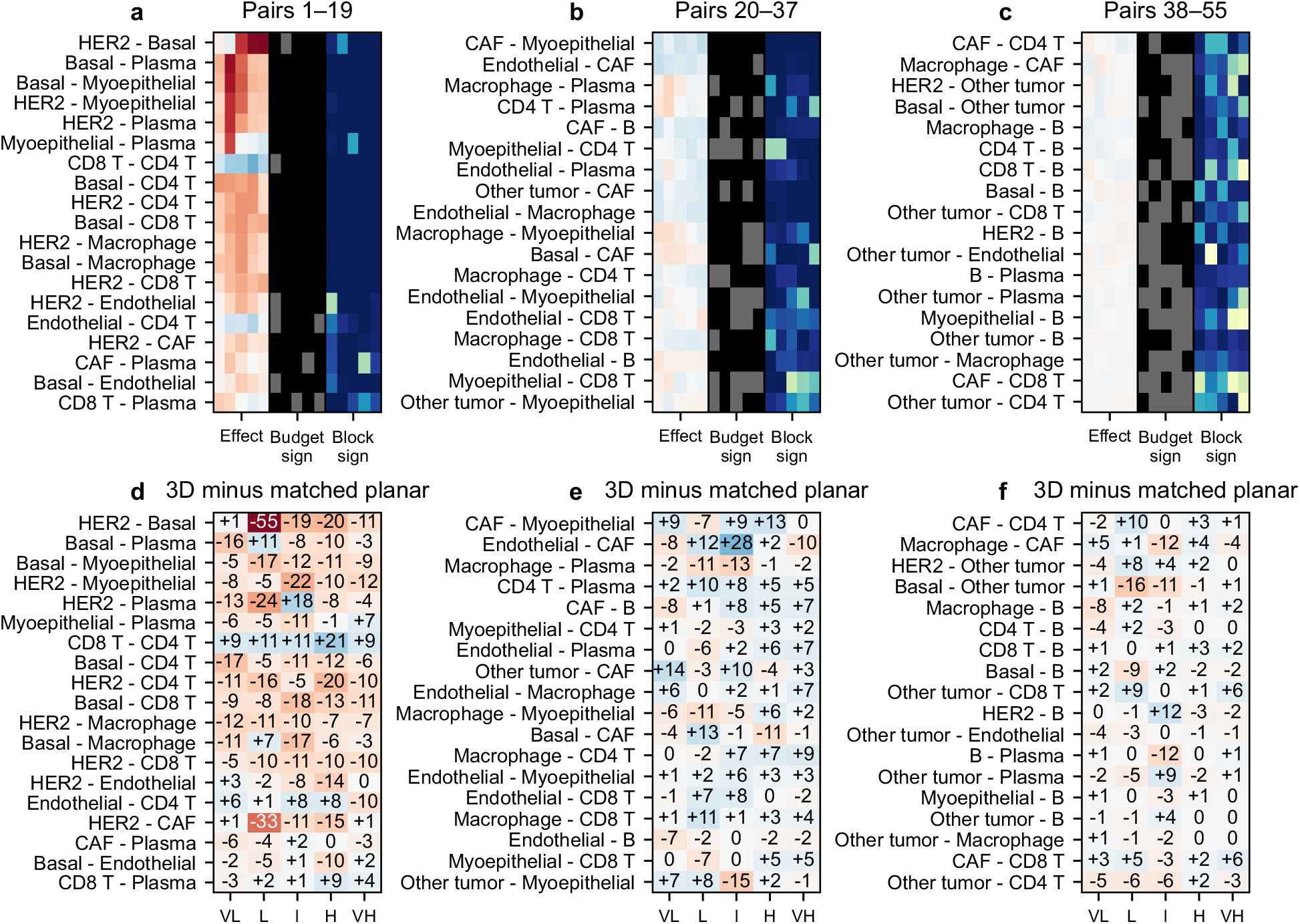
Complete breast quantitative comparison for all 55 active-variable pairs. Panels (**a**–**c**) show all active pairs across five relative panCK-burden strata. “Effect” is the signed fitted partial correlation; “Budget sign” is the fraction of three cell-sampling budgets matching the aggregate sign; and “Block sign” is the fraction of 100 serial-section block resamples retaining that sign with GP residuals fixed. Panels (**d**–**f**) show corresponding 3D-minus-matched-section-wise-2D partial correlations; cell numbers are differences in hundredths (e.g., +6 denotes approximately +0.06). Signed effect colors span −0.4 to +0.4; darker budget and block-sign cells represent stronger agreement. Stability measures are descriptive, and the lower panels compare complete estimators.

#### Zone-specific interaction patterns

In the Very Low breast zone, the largest positive fitted associations are Endothelial–CAF (0.093), CAF–Myoepithelial (0.083), and CD8^+^ T–CD4^+^ T (0.081); Basal tumor–CD4^+^ T is negative (−0.176). In Low, several tumor–stromal or tumor–plasma associations are more negative, including Basal tumor–Plasma (−0.349), Basal tumor–Myoepithelial (−0.330), and HER2-positive tumor–Plasma (−0.296). The HER2-positive–Basal tumor fitted effect is near zero and positive in Low (0.042), with only two of three sampling budgets retaining the aggregate sign, whereas it is negative in Intermediate (−0.290), High (−0.366), and Very High (−0.374), with three-budget sign agreement in those zones. This is a change in a conditional density summary, not evidence that one tumor subtype physically excludes another.

Across Intermediate, High, and Very High, CD8^+^ T–CD4^+^ T remains positive (0.143, 0.194, and 0.120). CAF–Myoepithelial is positive but modest in every zone (0.032–0.098); Endothelial–CAF is likewise modest (0.039–0.093). These cell pairs are biologically relevant to ductal and stromal architecture, but the fitted effects remain modest. Three-dimensional DCIS experiments document myoepithelial–CAF interplay^32^; other work links altered myoepithelial behavior to invasion^33,34^, and breast-tumor fibroblasts can support vascular growth^35,36^. These studies provide context for follow-up rather than validation of the fitted partial correlations. The lower overall budget-sign agreement in breast tissue and the one-specimen design require these zone patterns to remain exploratory.

The central matched 2D breast fit used the same 45,000 selected cells and the same downstream covariance estimator. The median absolute 3D-minus-2D difference among active pairs was 0.044, with a maximum of 0.549 for HER2-positive–basal tumor in Low (Figure 8). The planar comparator is sensitive to small section–zone groups and cannot be interpreted as a Z-only ablation. This large application contrast describes a change between complete spatial-adjustment strategies, not a uniquely volumetric biological association.

### Direct residual precision sensitivity

We fitted an EBIC-selected graphical lasso directly to the GP-adjusted residual matrix within each zone and sampling tier, with *γ* = 0.5 and a regularization grid from 0.005 to 0.5. Compared with the full MSFA zone covariance precision, Spearman correlations of pairwise partial correlations were 0.996–0.999 across CRC zones and 0.969–0.996 across breast zones (Appendix Figure 9); the sign of MSFA entries of magnitude at least 0.1 agreed in each zone. It says that the two precision estimators agree *conditional on these GP-adjusted residuals*; it does not cancel the much lower breast budget stability or resolve smooth-background ambiguity. The direct fit probes dependence on the MSFA covariance assumption, while budget and section-block checks probe different sensitivities.

**Fig A1.**
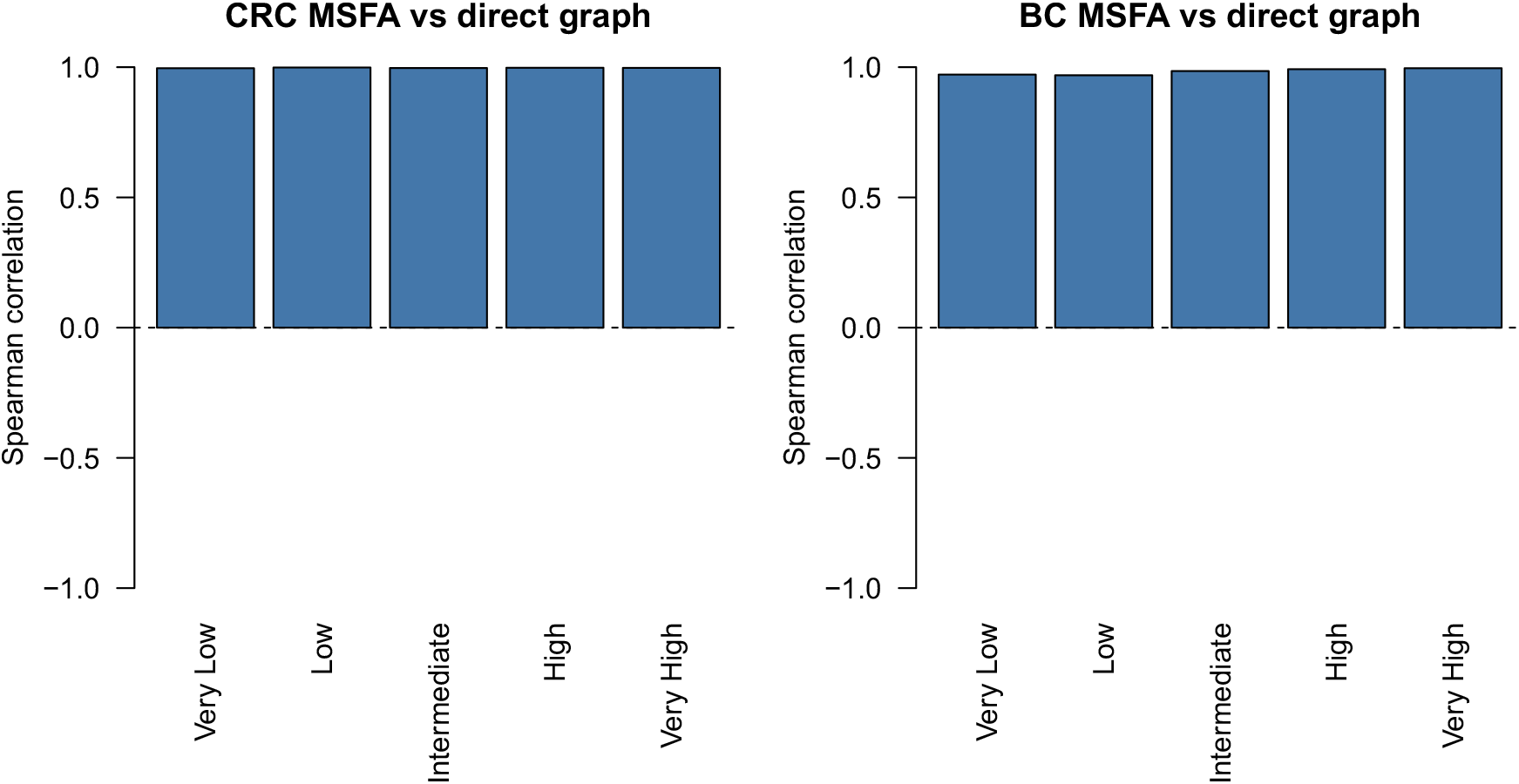
Direct residual-precision sensitivity. Spearman correlations between zone-specific MSFA and EBIC graphical-lasso partial correlations are shown for the five CRC and five breast zones. Both estimators use the same GP-adjusted residuals; the comparison does not establish biological replication or edge-wise uncertainty.

## Discussion

ISPat-3D integrates established GP and MSFA tools for a specific purpose: estimating shared and tumor-zone-varying conditional associations from registered 3D multiplex images. Its contribution is the integration, anisotropic 3D parameterization, and zone-resolved workflow, not a new GP, factor model, or universal 3D point-process method. Partial-correlation edges are descriptive conditional associations among measured density surfaces. They are not evidence of contact, signaling, direction, or mechanism.

The CRC analysis identifies positive conditional density associations within a T-cell set and between macrophage and stroma across this specimen’s relative tumor-density zones. Literature on CRC immune neighborhoods and fibroblast–macrophage niches makes these cell pairs biologically plausible candidates^3,30,31^. The fitted partial correlations do not show physical proximity of individual cells, immune function, or a mechanism. Their relatively high agreement across three sampling budgets and fixed-residual section-block resamples is more reassuring than the breast application, yet independent CRC volumes are needed to learn whether the pattern generalizes. In the breast specimen, HER2-positive–basal tumor associations differ by relative zone, while CAF–Myoepithelial and Endothelial–CAF associations are modest in the fitted analysis. These associations can guide imaging follow-up, but the 68.0% active-pair three-budget sign agreement makes rankings fragile. The strong direct graphical-lasso agreement on the *same* GP-adjusted residuals checks one covariance-estimation choice; it is compatible with instability from cell selection, small section–zone groups, registration, and GP scale. Breast conclusions remain strictly exploratory.

The Myoepithelial ↔ CAF candidate has direct experimental precedent: a three-dimensional DCIS model demonstrated functional interplay between human myoepithelial cells and CAFs^32^, while other studies linked altered myoepithelial behavior to invasive progression^33,34^. The CAF ↔ Endothelial candidate is also biologically plausible because breast-tumor fibroblasts can promote endothelial and vascular growth in experimental systems^35,36^. Nearest-neighbor, neighborhood-composition, point-process, graph, and GP variance-component methods answer related but different questions^13,14,19^. ISPat-3D is useful when the target is a conditional-association covariance decomposition across ordered volumetric zones. The anisotropic GP accommodates unequal in-plane and axial scales; MSFA borrows information across zones; and the precision transformation gives a compact conditional summary. These components can be used separately, as the ablation and residual-graph analyses illustrate. Claims beyond registered 3D multiplex imaging remain prospective.

The key integration is registered three-coordinate GP adjustment followed by shared and zone-varying factor covariance and conditional-association summaries. Vecchia neighbors and spatially balanced anchors make GP fitting feasible at the reported cell budgets^37,38^. The Gaussian-likelihood factor fit uses full residual covariance summaries and exact zone counts while preserving the shared-plus-zone covariance form. The planar comparison follows the same stages on identical cell rosters, with GP fits made independently by section. Its application deltas contrast two complete spatial-adjustment strategies, so they do not assign a difference to *z* alone. The focused controlled experiment demonstrates a benefit of axial information only under its known depth-varying nuisance design.

The GP separates patterns by modeled smoothness, not by biological cause. Smooth tumor fronts, stromal boundaries, vascular organization, or immune exclusion may be removed together with technical gradients. Conversely, residuals can carry fine-scale spatial structure, noise, segmentation or annotation error, and effects of unmeasured cell populations. The short-scale GP experiment shows that reducing Moran autocorrelation need not improve edge recovery. No edge intervals, posterior edge probabilities, or independent-specimen replication are provided; budget and fixed-residual block sign stability must not be read as significance. The five density zones are relative within each image, and their boundaries need not map to discrete tissue states.

Registration uncertainty is a priority for future benchmarking. Even modest XY displacement among sections can change inferred axial correlation and a pooled volumetric GP; unequal section spacing and errors in segmentation or cell typing can also matter. A larger registered cohort, perturbations of registration and zone boundaries, and comparisons with explicit spatial point-process or interaction assays would be needed to distinguish technical sensitivity from reproducible biology. The present work establishes a feasible, transparent analysis of one CRC and one breast volume and identifies biologically plausible, tumor-zone-resolved conditional associations for further testing.

## Methods

### Simulation study

#### Data-generating mechanism

The primary simulation tested recovery of shared and zone-specific covariance after removal of a spatial nuisance. Each realization contained *Q* = 3 zones. Within zone *q*, planar coordinates were drawn independently from normal distributions centered at (3*q*, 3*q*) with standard deviation 1.5 in each direction, and depth was drawn from Uniform(0, 1). This gives partially overlapping planar regions and a common depth range in arbitrary simulation units, rather than a particular tumor geometry. For *G* variables, the shared factor count was *K* = ⌈ 2log *G*⌉. Entries of the *G* × *k* shared loading matrix ***Φ*** were drawn from *N*(0, 1). For each zone, the number of generating zone-specific factors was sampled independently and uniformly from {2, …, *K* − 1}; its loading matrix ***Λ***_*q*_ also had independent *N*(0, 1) entries. Diagonal uniqueness values were independent Uniform(0.1, 1) draws. The generating covariances were

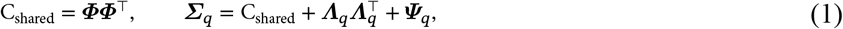

and the nonspatial cell profiles were independent Z_*q*_ ~ *N* _*G*_(0, ***Σ***_*q*_) within each zone. The covariance was therefore generally dense, despite its low-rank-plus-diagonal construction.

For each variable and zone, an independent zero-mean spatial field *B*_*g*_ had covariance 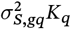, with 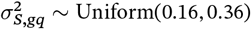. The generating *K*_*q*_ was either an anisotropic Matérn-3/2 or radial-basis-function (RBF) kernel. Both used a common planar range *l*_*S*_ for *x* and *y* and a separate axial range *l*_*Z*_ for *z*. The two ranges were the medians of the pooled positive within-zone pairwise planar and axial distances, respectively, across the three zones; they were not recalculated separately for each zone. With *β*_*g*_ ~ Uniform(1, 3) and independent *ϵ*_*q*_ ~ *N*^2^), the observed response was

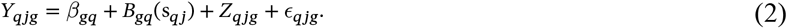

The spatial fields and nonspatial factor signal were independent by construction. Thus the known C_shared_ and ***Σ***_*q*_ provide targets for the two-stage recovery assessment.

### Design and evaluation

We evaluated *G* ∈ {15, 20, 25} and six zone-size vectors (*N*_1_, *N*_2_, *N*_3_): (300, 300, 300), (250, 250, 250), (250, 300, 300), (250, 250, 300), (300, 250, 250), and (300, 300, 250). Matérn-3/2 and RBF generating kernels each had five independent replications per size and *G*, giving 180 data sets. The analysis fit a Matérn-3/2 spatial GP to each variable within each zone using the 15-neighbor Vecchia approximation and then fitted the same Gaussian-likelihood shared-plus-zone covariance form used for the applications. Its shared and zone-specific fitted loading ranks were both ⌈2 log *G*⌉; the generating zone-specific ranks varied as described above. The RBF settings therefore also test recovery under a different generating kernel. The shared target was C_shared_, whereas each zone-level target was the *full* ***Σ***_*q*_, including uniqueness. We compared each fitted matrix with its corresponding target using the RV coefficient^28^.

For a second descriptive metric, off-diagonal entries with absolute covariance greater than 0.1 were labeled above threshold in truth and fit. Sensitivity was the proportion of above-threshold true entries retained by the fit. The reported false-discovery proportion was the fraction of fitted above-threshold entries whose true magnitude did not exceed 0.1. Since normally drawn loadings yield almost surely dense covariance, this threshold distinguishes magnitudes rather than identifying structural zero edges or controlling an inferential false-discovery rate. The value 0.1 was fixed for simulation evaluation and differs from the adaptive display cutoffs used to reduce edge crossings in application network drawings. GP-stage and covariance-fit wall times were recorded in the observed computational environment.

### Focused GP deconfounding validation

The primary factor simulation generates independent spatial fields for different variables. To test the GP where a *shared* smooth nuisance can itself create cross-variable dependence, we used *N* = 300 locations and *G* = 12 variables. Planar coordinates were independent Uniform(0, 1) draws and depth was Uniform(0, 0.4). The nonspatial signal had independent rows Z_*j*_ ~ *N* _*G*_(0, ***Σ***), where ***Ω*** = ***Σ***^−1^ had diagonal entries 1, entries −0.25 between adjacent nodes of a 12-variable ring, and zero elsewhere. Thus adjacent true partial correlations were 0.25 and all other off-diagonal partial correlations were exactly zero. The observed matrix was

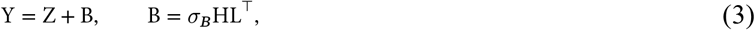

where the two columns of H were independent Gaussian fields with the same anisotropic generating kernel *K*_0_, and row *g* of L was (cos *a*_*g*_, sin *a*_*g*_) for equally spaced angles *a*_*g*_. Each row of L has unit norm, so every variable has nuisance variance 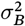, while the two shared fields induce cross-variable spatial confounding. Generating ranges were *l*_*S*_ = 0.25 and *l*_*Z*_ = 0.12 for both Matérn-3/2 and RBF kernels. We varied *σ*_*B*_ ∈ {0, 0.5, 1.0, 1.5} and generated 20 independent realizations per kernel and strength.

Four adjustments were applied to the *same* generated realization in each replicate. The matched 3D arm used the generating kernel family and both generating ranges. The short-scale 3D arm halved the planar and axial ranges to 0.125 and 0.06. The planar arm used the generating kernel family and planar range but only (*x, y*) coordinates. The no-GP arm subtracted the column mean alone. For each GP arm, a 15-neighbor Vecchia approximation predicted the spatial field at the observed locations using the known nuisance variance and the average nonspatial signal variance *τ*^2^ = *G*^−1^ tr(***Σ***). Its residual was 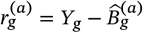 for arm *a*. These parameters were fixed at their generating values, except for the deliberately altered ranges and omitted depth in the corresponding arms, to isolate the adjustment step. This is a component validation, not a test of estimated GP hyperparameters or the complete factor-fitting pipeline. At *σ*_*B*_ = 0, all four arms reduce to mean-centering.

We evaluated (i) mean absolute deviation of eight-nearest-neighbor Moran’s *I* from its randomization expectation −1/(*N* − 1), (ii) root-mean-square error of all upper-triangle estimated partial correlations relative to the ring truth, (iii) the proportion of exactly null pairs with 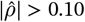, and (iv) root-mean-square error of the adjusted residuals relative to the centered nonspatial signal. Lower values are preferable on all four metrics. The fixed 0.10 rule in (iii) is a diagnostic cutoff, not a significance threshold.

### Inputs and relative tumor-density zones

Each sampled location has registered coordinates s_*j*_ = (*x*_*j*_, *y*_*j*_, *z*_*j*_), a section identifier, a specimen-specific zone label *q*_*j*_ ∈ {1, …, 5}, and cell-type neighborhood-density estimates (KDEs). The published CRC CyCIF and breast IMC processing and annotation protocols are described in Appendices A G and A H^8,9^. We use log(1 + 10^9^KDE) for each cell-type variable. Five ordered strata are derived from each specimen’s relative tumor-burden proxy: a smoothed tumor-cell KDE surface in CRC and panCK intensity in breast. The labels run Very Low to Very High. Tied breast scores yield unequal zone populations, and adjacent boundaries have no asserted discrete biological meaning.

CRC source tables contained 9,431,098 processed rows across 25 sections. Sampling budgets were 50,000, 100,000, and 150,000 *per zone*, for total selected rosters of 250,000, 500,000, and 750,000 cells. Breast source tables contained 73,936 cells across 152 sections. Its budgets were 25,000, 45,000, and 65,000 *total across zones*; zone allocations were proportional to available counts. Within each zone, section allocations were proportional to section population. Selection was spatially balanced over a 10-by-10 in-plane grid within each section, with a fixed tier seed, and source row identifiers were saved. These rosters are the actual inputs to all fitted analyses and figures.

### Spatial GP adjustment

For cell-type variable *g* in zone *q*, the working model separates an intercept, a smooth spatial field, and a residual:

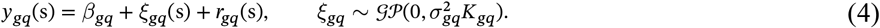

The three-coordinate Matérn-3/2 kernel is anisotropic^18,39^. Its scaled distance is *d*^2^ = (*Δ*x*x*)^2^ + (*Δ*y*x*)^2^ + (*Δ*z*x*)^2^; distinct positive axial and in-plane ranges allow a registered depth gradient to differ from a planar gradient. This is motivated by the genuine three-coordinate structure of these data and by the general need to represent anisotropy, rather than by a claim that depth alone causes any estimated edge. The RBF kernel was also used as a generating alternative in simulation; the applications used Matérn-3/2.

For large fits, we used GPBoost’s 15-neighbor Vecchia likelihood approximation^37,38^. The likelihood factors each anchor response conditionally on a small set of nearby predecessors, avoiding a dense covariance factorization. For each zone and variable, anchors were selected spatially across sections from the selected roster: approximately 10% of selected cells, with a minimum of 300 and a maximum of 5,000 anchors. Positive covariance parameters and the intercept were optimized with a maximum of 30 iterations; fitted fields were predicted at *all* selected cells and subtracted, 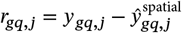. Constant variables were mean-centered. Thus only GP parameter fitting used anchors; every selected row informed the downstream residual covariance. The Vecchia approximation avoids the cubic time and quadratic storage of a dense GP covariance factorization.

This adjustment operationally defines broad modeled variation as background. It does not identify whether that smooth variation is technical or biological. The range parameters, anchor selection, and section registration can change the residuals, and unmodeled structure may remain^27^. The focused simulation examined half-scale and planar alternatives under a known nuisance field; it does not remove this ambiguity from the applications.

### Shared and zone-specific factor covariance

The adjusted *N*_*q*_ × *G* residual matrix in zone *q* yields its complete sample covariance S_*q*_ and actual row count *N*_*q*_. We use the MSFA covariance form^23,24^:

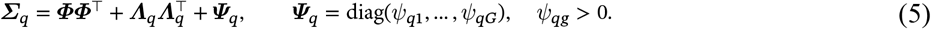

Here ***Φ***^⊤^ is covariance shared across zones and 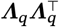 is a zone-specific covariance contribution. Both loading ranks were five for the CRC and breast applications. The breast luminal tumor KDE was identically zero and was omitted from the active likelihood, then re-embedded as a zero row and column for label provenance. Breast effect and delta figures report only pairs among the 11 active variables. In simulations the rank was ⌈2 log *G*⌉.

The large-data fit is a Gaussian covariance maximum-likelihood fit from the sufficient statistics {(*N*_*q*_, *S*_*q*_)}. With 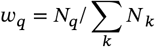, its loss, up to constants, is

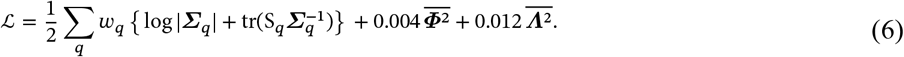

Each variable was scaled by its pooled residual standard deviation before optimization and rescaled afterward. Positive uniqueness values used a softplus parameterization; the shared and zone loadings were initialized from positive eigencomponents and optimized by L-BFGS in double precision^40,41^. The loading matrices are rotationally non-identifiable, so we interpret their covariance products and the full fitted zone covariance, not individual loading columns. The likelihood targets the stated covariance form without a variational posterior interpretation.

### Conditional association and stability

The zone precision matrix is the inverse of the *full* fitted covariance, with a 10^−6^ numerical diagonal ridge if needed. For distinct variables *g* and *h*,

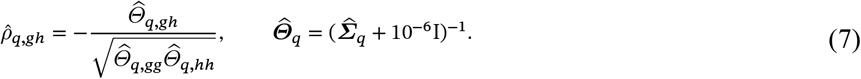

A signed edge means conditional association among adjusted measured density variables under this working covariance model. The shared loading product is low rank and has no unregularized inverse; all reported zone edges use the full covariance including zone loadings and uniqueness.

For each pair and zone, we summarized the Fisher-transformed average partial correlation across three sampling budgets and counted how many tier estimates retained the aggregate sign. We also made 100 serial-section block resamples within the central-tier data using *fixed* GP residuals and section-level covariance sufficient statistics, refit the covariance precision, and recorded the fraction of resamples retaining each sign. These are stability descriptions without GP refitting, registration perturbation, posterior edge probabilities, or confidence intervals. They cannot support inferential significance.

### Matched serial-section planar comparison and direct precision sensitivity

The planar comparator follows the published ISPAT architecture^26^: cell-type KDE, GP spatial adjustment, shared-plus-zone covariance, and conditional-association summary. We implemented the same stages with a scalable likelihood for these cell rosters. It uses precisely the selected row identifiers from the central 3D tier: 100,000 CRC cells per zone or 45,000 breast cells total. A Matérn-3/2 Vecchia GP is fit separately to (*x, y*) within each section–zone group; the full set of planar residuals then enters the same covariance-likelihood MSFA fit. Groups with fewer than ten cells and constant variables are mean-centered. The reported 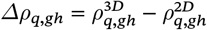 compares pooled volumetric spatial fitting against independent section-wise planar spatial fitting on identical cells. The comparison is matched in cell roster and downstream estimator, but changes both the coordinate dimension and section pooling; it is not a Z-only causal ablation.

We also fit sparse precision matrices directly to each zone’s GP residuals by graphical lasso. The regularization parameter was chosen by extended BIC with *γ* = 0.5 over 40 values from 0.005 to 0.5 after variable scaling. We compared its signed partial correlations with those derived from full MSFA zone covariances. This probes sensitivity to the factor covariance form conditional on the same residuals and does not assess independent biological replication.

### Optional reference-informed covariance regularization

External knowledge can optionally be used to reweight the MSFA covariance components. Let A ∈ ℝ^*G* × *G*^ be a symmetric, nonnegative adjacency or similarity matrix defined on exactly the same ordered cell-type labels as the fitted model. Such a reference may summarize curated cell–cell relationships, receptor–ligand-supported cell-type pairs, or an independently constructed spatial-proximity graph. Because an adjacency matrix need not be positive semidefinite, it cannot in general be inserted directly as a Gaussian covariance. We first symmetrize it and construct a positive-semidefinite reference covariance ***Σ***^∗^, either by projecting A to the nearest positive-semidefinite matrix^42^ or by eigendecomposition, replacing negative eigenvalues by zero, and adding a small diagonal ridge when positive definiteness is required. Rescaling ***Σ***^∗^ to unit diagonal prevents arbitrary reference magnitude from determining the overall covariance scale.

Let [***Σ***^∗^]^1/2^ = U***Δ***^1/2^U^⊤^ denote its symmetric square root. The reference-weighted shared and zone-specific factor covariances are

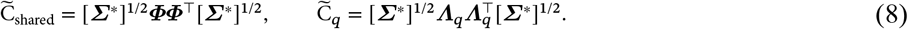

Both matrices remain positive semidefinite. For example, 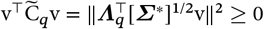. The complete reference-informed covariance passed to precision estimation is

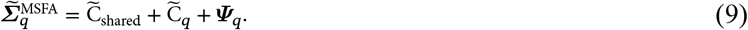

This construction is a soft covariance-scale regularization: it emphasizes directions supported by the external reference while retaining the data-estimated shared and zone-specific loading products. It is not a hard constraint that forces a referenced pair to appear as a partial-correlation edge, nor does it turn an adjacency entry into evidence of signaling in the analyzed tissue. Reference construction must therefore be prespecified, label-matched, and kept independent of the outcomes used to evaluate the network. When no credible external reference is available, setting ***Σ***^∗^ = I_*G*_ recovers the unregularized full covariance 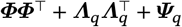. The CRC and breast-carcinoma analyses reported here used this identity-reference setting; the optional construction is provided for future reference-informed applications. The output for one image is one regularized shared-factor component summary and *Q* partial-correlation graphs obtained from the full fitted zone covariances. They summarize measured association structure and do not identify ligand–receptor pairs or causal effects.

### Generative AI use

OpenAI GPT-5.2 (OpenAI, 2025) was used to refine manuscript language and tone, format BibTeX citations, improve LaTeX displays (including algorithms and tables), and assist with R analysis and figure-generation scripts. GPT-5.2 and Claude Sonnet 4.6 (Anthropic, 2026) were also used to annotate, streamline, and document the R package for public release on GitHub. The authors reviewed and revised all AI-assisted text, code, and displays, checked the resulting outputs, and retain full responsibility for the methods, results, and conclusions.

## Detailed network summaries

The complete pair-by-zone fitted effects and descriptive tier sign stability are shown without edge filtering in Figures 6 and 8. The complete 3D-minus-matched-section-wise-2D differences are shown in Figures 6 and 8. Under the working Gaussian model, these are conditional associations among measured KDE variables after GP adjustment; they do not identify physical contacts or mechanisms. The shared factor covariance is low rank and has no separate unregularized precision matrix, so only complete zone covariances support the reported partial correlations.

The CRC pair–zone sign agreement across the three actual budgets was 169/180, versus 187/275 among active breast pairs. The corresponding fractions with at least 90% sign agreement in fixed-residual serial-section block resamples were 87.2% and 62.2%. The section resampling does not refit spatial GPs or perturb registration and must not be interpreted as an edge-wise confidence interval. The effect tables and their production scripts are described in the released code workflow.

### Direct residual graphical-lasso sensitivity

The graphical-lasso comparison uses the same GP-adjusted residuals as the large-fit analysis. As described in Results, its strong agreement with the factor-covariance precision is conditional on these residuals and does not measure replication or uncertainty from registration, annotation, or GP fitting See fig A1.

### CRC data processing

#### Acquisition and column standardization

Per-section cell feature tables were downloaded for all 25 CRC1 serial sections. Each feature table was standardized to a common schema. Global tissue-level coordinates (*X*_*s*_, *Y*_*s*_) from the registered whole-slide images were used directly as the canonical XY position for each cell, ensuring spatial consistency across sections. A panel of 23 markers was retained: epithelial markers (Keratin, E-cadherin, CDX2), immune lineage markers (CD45, CD3, CD4, CD8a, FOXP3, CD20, CD68, CD163, CD45RO), proliferation markers (Ki67, PCNA), stromal markers (*α*-SMA, Vimentin, Desmin, Collagen, CD31), checkpoint markers (PDL1, PD1), and structural markers (NaKATPase, LaminABC). Marker intensity values were winsorized at the 99.9th percentile to mitigate the influence of imaging outliers. Cells with missing XY coordinates or any missing marker intensity after winsorization were excluded, as were sections with fewer than 1000 retained cells.

#### Cell type classification

Cell types were assigned by hierarchical threshold gates applied independently to each section, following^9^. Nine mutually exclusive labels were defined in this order: tumor cells (Keratin > 7000, E-cadherin > 3500, or CDX2 > 15000), Treg (FOXP3 > 200 and CD4 > 1900), CD8^+^ T (CD8a > 2500), CD4^+^ T (CD4 > 1900), other T (CD3 > 5800), B cells (CD20 > 3300), macrophages (CD163 > 760 or CD68 > 430), other immune cells (CD45 > 2000), and stromal cells (*α*-SMA > 3300, Vimentin > 560, or Collagen > 7700). The first satisfied gate determines the label. Thus “other T” is the residual CD3^+^ category after Treg, CD8^+^, and CD4^+^ assignments and does not overlap them. Unclassified cells were excluded.

#### Spatial zone annotation via tumor cell density partitioning

Tumor cell spatial density serves as a biologically meaningful stratification axis for partitioning each tissue section into spatially coherent microenvironmental compartments. Tumor cell density varies systematically across a tissue section, encoding the spatial gradient from immune-rich, tumor-sparse stromal and peri-tumoral regions to dense tumor epithelial cores. This gradient is particularly informative in the context of CRC, where the spatial organization of immune and tumor cell compartments differs substantially across the invasive margin morphologies documented in CRC1. We therefore estimate a spatially smoothed tumor cell density surface via two-dimensional kernel density estimation (KDE) and use it as the stratification axis to partition each section into relative spatial strata prior to downstream inference.

For each section, the XY coordinates of all classified tumor cells were extracted and a Gaussian KDE surface was fitted on a 200 × 200 grid spanning the full XY extent of the section. The kernel bandwidth was selected separately along each spatial axis by leave-one-out cross-validation (LOOCV). Specifically, for a univariate coordinate vector x = (*x*_1_, …, *x*_*n*_), the LOOCV objective is:

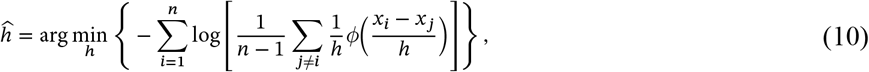

where *ϕ*(⋅) denotes the standard Gaussian density. The search interval for *h* was set to [0.05 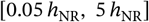, where 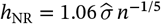 is the Gaussian normal-reference bandwidth^43,44^, and optimization was carried out using Brent’s method^45^. When the number of tumor cells in a section exceeded 5000, a random subsample of 5000 cells was used for bandwidth estimation while the full tumor cell set was retained for KDE fitting. Sections with fewer than 50 tumor cells were assigned uniformly to zone 1.

The fitted KDE surface was bilinearly interpolated at the XY location of every cell *j* in the section, regardless of cell type, yielding a continuous local tumor density score 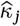. For each tissue section ℐ, let 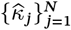 denote these scores across all *N* cells at spatial locations {*s*_*j*_}. We partition ℐ into five non-overlapping zones 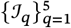 by thresholding on the empirical quantiles of 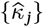:

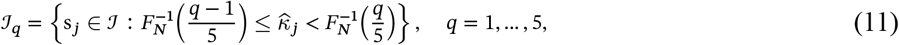

where 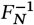 is the empirical quantile function of 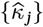 across all *N* cells, and the closed upper bound in the final stratum ensures no cell is excluded at the maximum density value. The five zones are labeled Very Low (*q* = 1, 0–20th percentile), Low (*q* = 2, 20–40th percentile), Intermediate (*q* = 3, 40–60th percentile), High (*q* = 4, 60–80th percentile), and Very High (*q* = 5, 80–100th percentile), corresponding to increasing local tumor cell density. Zone 1 therefore represents the tumor-sparse periphery and zone 5 the tumor-dense core. By construction the zones are mutually exclusive and exhaustive: 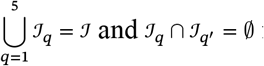for q≠q′.

Quintiles adapt to section-level density shifts and provide approximately balanced strata for estimation. Five zones were selected to retain the five-level tumor-gradient convention of ISPAT while avoiding very small strata; this is a pragmatic resolution choice rather than a uniquely biological number. The zones, not the serial sections, are the *Q* = 5 study-specific inputs to MSFA; registered sections jointly supply the 3D coordinates within each zone.

#### Cell type-specific KDE surface construction

To obtain a continuous, spatially resolved compositional representation of the TME at each cell location, we computed cell type-specific KDE surfaces for all nine cell types across each of the 25 CRC1 sections. For each section and each cell type *t* ∈ *T*, the XY coordinates of cells assigned to type *t* were used to fit a separate two-dimensional Gaussian KDE surface on a 200 × 200 grid spanning the section’s full spatial extent, using the same LOOCV bandwidth selection procedure described in Equation (10). Cell types with fewer than 30 cells in a given section were assigned a density of zero throughout that section. The fitted surfaces were bilinearly interpolated at the XY coordinates of every cell in the section, yielding a per-spatial location vector of local neighborhood densities:

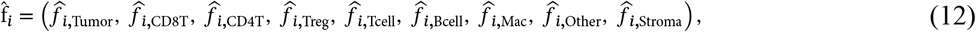

for spatial location *i*. This encodes the local cellular composition of each cell’s spatial neighborhood in a continuous and spatially smooth form, and serves as the input variable matrix to the ISPat-3D conditional-association pipeline.

### BC data processing

#### Acquisition and cell centroid extraction

Per-section single-cell mean intensity tables were downloaded from the Zenodo repository. Cell centroids in three dimensions were extracted from the 3D segmentation mask yielding physical coordinates (*X, Y, Z*) in micrometres at 1 *µ*m/pixel resolution in the XY plane and 2 *µ*m per section along the Z axis. Centroid tables were joined to the mean intensity table on cell label. Marker intensity values were winsorized at the 99th percentile to mitigate the influence of imaging outliers.

#### Cell type classification

Cell types were assigned using a hierarchical threshold-based gating strategy applied to 99th-percentile normalized marker intensities, analogous to the classification framework employed for the CRC1 specimen. Twelve cell types were defined in the following order of precedence: endothelial cells (vWF/CD31 > 0.3), macrophages (CD68 > 0.3), B cells (CD20 > 0.3 and CD45 > 0.2), plasma cells (CD138 > 0.3), CD8^+^ cytotoxic T cells (CD3 > 0.3 and CD8a > 0.2), CD4^+^ helper T cells (CD3 > 0.3 and CD8a ≤ 0.2), cancer-associated fibroblasts (CAF; Collagen I > 0.3, SMA > 0.3, and panCK < 0.2), myoepithelial cells (SMA > 0.3 and CK5 > 0.3 or CK14 > 0.3), and four epithelial tumor subtypes distinguished by their cytokeratin expression profiles: HER2-positive tumor cells (panCK > 0.3 and HER2 > 0.4), basal tumor cells (panCK > 0.3 and CK5 > 0.3 or CK14 > 0.3), luminal tumor cells (panCK > 0.3 and CK7 > 0.3), and other tumor cells (panCK > 0.3). Labels were assigned in the listed order of precedence. Cells satisfying no criterion were labeled unassigned and excluded from all downstream analyses. *Spatial zone annotation via pan-cytokeratin intensity partitioning* Unlike the CRC1 analysis, where spatial zones were defined by a KDE-smoothed tumor cell density surface, the compact single-tissue-block geometry of the breast carcinoma dataset and the direct availability of per-cell pan-cytokeratin (panCK) intensities permit a simpler and equally principled stratification strategy. Pan-cytokeratin is a broad epithelial lineage marker that captures all cytokeratin-expressing tumor cells regardless of molecular subtype and therefore serves as a continuous proxy for local tumor epithelial burden at each cell location.

The raw panCK intensity was normalized to the 99th percentile of the global distribution, yielding a normalized score 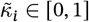. The five spatial zones were then defined by partitioning the global empirical distribution of 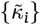 at its quintiles, following the same quantile-based stratification used for CRC1 in Equation (11). The five zones are labeled Very Low, Low, Intermediate, High, and Very High, corresponding to increasing local panCK expression as a relative epithelial-burden proxy in this specimen. This approach is directly comparable to the KDE-based stratification used for CRC1 in that both methods partition cells along a continuous, specimen-specific tumor-burden proxy into five ordered relative strata; tied intensity scores make their observed cell counts unequal. Using the measured panCK intensity avoids a second smoothing step for the zone-defining score.

#### Cell type-specific KDE surface construction

To obtain a continuous, spatially resolved compositional representation of the TME at each cell location, we computed cell type-specific KDE surfaces for all 12 cell types across each of the 152 sections, following the same procedure applied to the CRC1 specimen. For each section and each cell type *t* ∈ *T*, the XY coordinates of cells assigned to type *t* were used to fit a separate two-dimensional Gaussian KDE surface on a 200 × 200 grid spanning the section’s full spatial extent, using the LOOCV bandwidth selection procedure described in Equation (10). The fitted surfaces were bilinearly interpolated at the XY coordinates of every cell in the section, yielding a per-location vector of local neighborhood densities 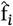 for spatial location *i*. This encodes the local cellular composition of each cell’s spatial neighborhood in a continuous and spatially smooth form, and serves as the input variable matrix to the ISPat-3D conditional-association pipeline.

## Author contributions

S.B. conceived the statistical methodology, implemented the computational framework, performed all analyses, generated all figures, and drafted the manuscript. A.R. contributed to study conception and critically reviewed the manuscript. All authors reviewed and approved the final version.

## Data availability

The CRC1 CyCIF dataset is publicly available from the Lin et al. (2023) data release. The 3D IMC breast carcinoma dataset is available from Zenodo (https://doi.org/10.5281/zenodo.4752030).

## Code availability

The ISPAT3D R package and Vecchia-GP analysis scripts are available at https://github.com/sagnikbhadury/ISPAT-3D (version 0.3.0: https://github.com/sagnikbhadury/ISPAT-3D/releases/tag/v0.3.0). The archived version-sp ecific software DOI is https://doi.org/10.5281/zenodo.22798429. The release README specifies processed-table inputs, actual per-zone or total-cell budgets, likelihood specification, simulations, matched section-wise 2D fitting, graphical-lasso sensitivity, and figure regeneration.

## Declarations

### Open Access

This article is licensed under a Creative Commons Attribution 4.0 International License, which permits use, sharing, adaptation, distribution and reproduction in any medium or format, as long as you give appropriate credit to the original author(s) and the source, provide a link to the Creative Commons licence, and indicate if changes were made. The images or other third party material in this article are included in the article’s Creative Commons licence, unless indicated otherwise in a credit line to the material. If material is not included in the article’s Creative Commons licence and your intended use is not permitted by statutory regulation or exceeds the permitted use, you will need to obtain permission directly from the copyright holder. To view a copy of this licence, visit http://creativecommons.org/licenses/by/4.0/.

